## Supplementary Figures 1-7 and Supplementary Table 1 for "Concerted localization-resets precede YAP-dependent transcription"

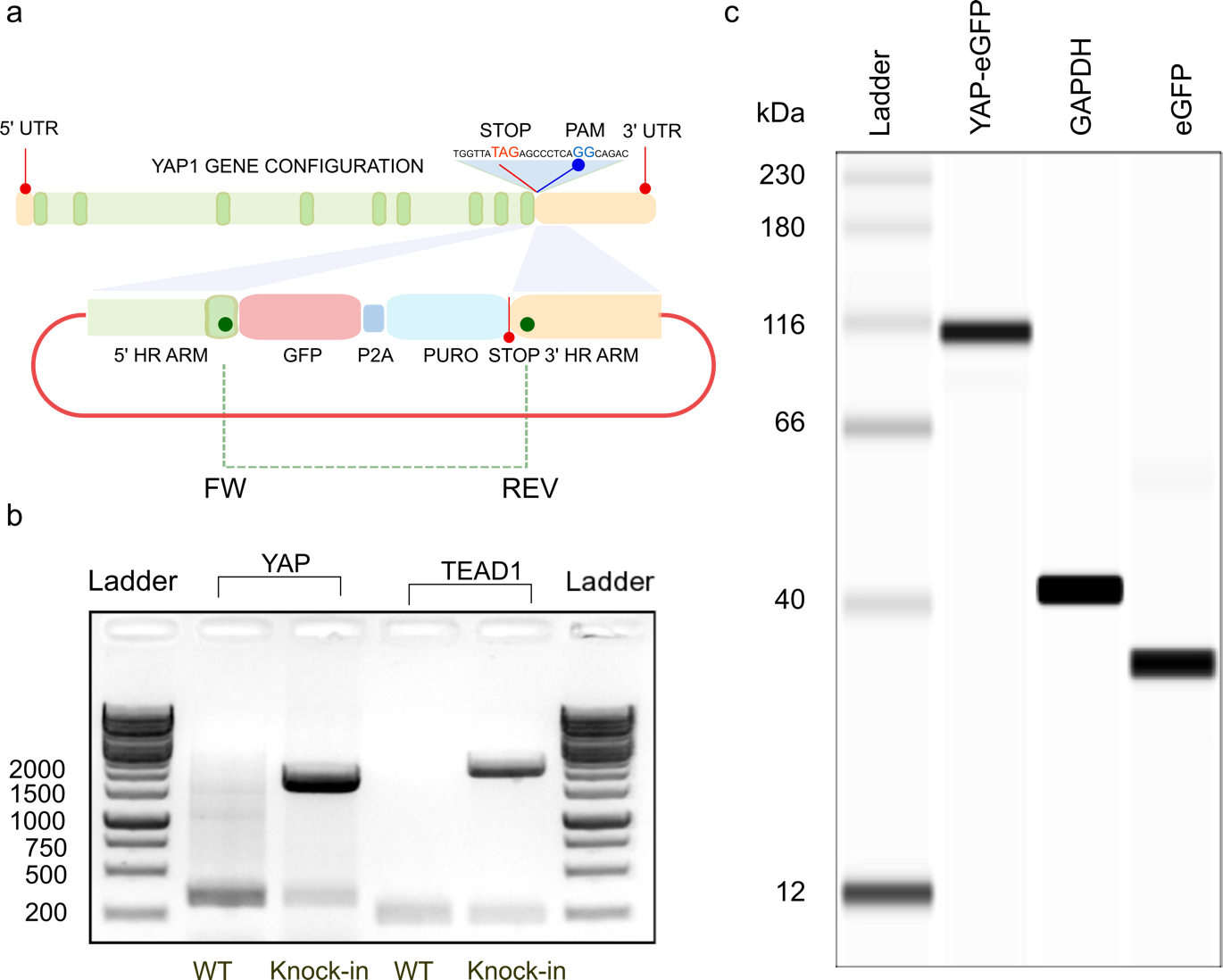

**Supplementary Fig. 1: Validation of Fluorescent Protein knockin cell lines. a.** Top: Cartoon of CRISPR-Cas9 based insertion of eGFP-P2A-puromycin cassette at 3’ end of the YAP gene as shown in **main text Figure 1a**. Two sites in the 5’ and 3’ HR arms are marked to show locations of the FW and Rev primers used for PCR based validation of knock-in. Similar primer design was used to validate mCherry-P2A-Hygromycin cassette knockin at the 3’ end of the TEAD1 gene.

**b.** Agarose gel electrophoresis of amplicons produced by PCR using YAP (lanes 2, 3) and TEAD1 (lanes 4, 5) primers sets against total genomic DNA extracted from WT (wildtype) and KI (knock-in) MCF10A cells. **c.** Capillary western blot using anti eGFP (lane 2) and anti GAPDH (lane 3) antibodies against MCF10A^YAP-GFP-KI^ whole cell lysate, and anti eGFP antibody against recombinant eGFP (lane 4). Observed vs expected (theoretical) sizes: Yap-eGFP (110/100), GAPDH (43/36), eGFP (35/29).

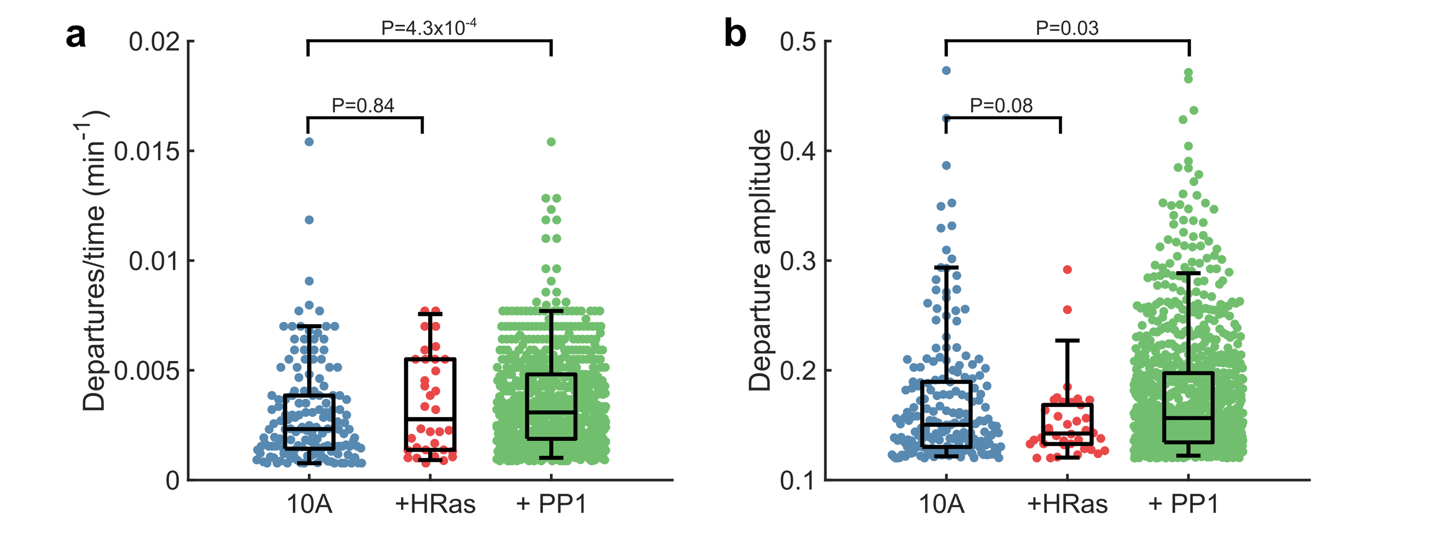

**Supplementary Fig. 2.** YAP N/C pulse density and amplitude analysis under different conditions (10A, 10A+HRas, 10A+PP1). Box and whisker lines: black-center line is median value, box-bounds are inter-quartile range, and whiskers are 0.05 and 0.95 quantiles. Data compiled from two independent 24-hr experiments for each condition. P-values are from two-sided Mann-Whitney U-test to test whether fluctuation frequency or amplitude is changed from MCF10A. (**a**) *Frequency*: N_10A_=150, N_HRas_=34, N_PP1_=693 tracks. 10A vs +HRas (P=0.84, Ranksum=13596, Z-val=-0.99), 10A vs +PP1 (P=4.33x10^-4^, Ranksum=54295, Z-val=-3.33 ). (**b**) *Amplitude*: N_10A_=192, N_HRas_=38, N_PP1_=970. 10A vs +HRas (P=0.08, Ranksum=22683, Z-val=1.35), 10A vs +PP1 (P=0.03, Ranksum=103896, Z-val=-1.8245).

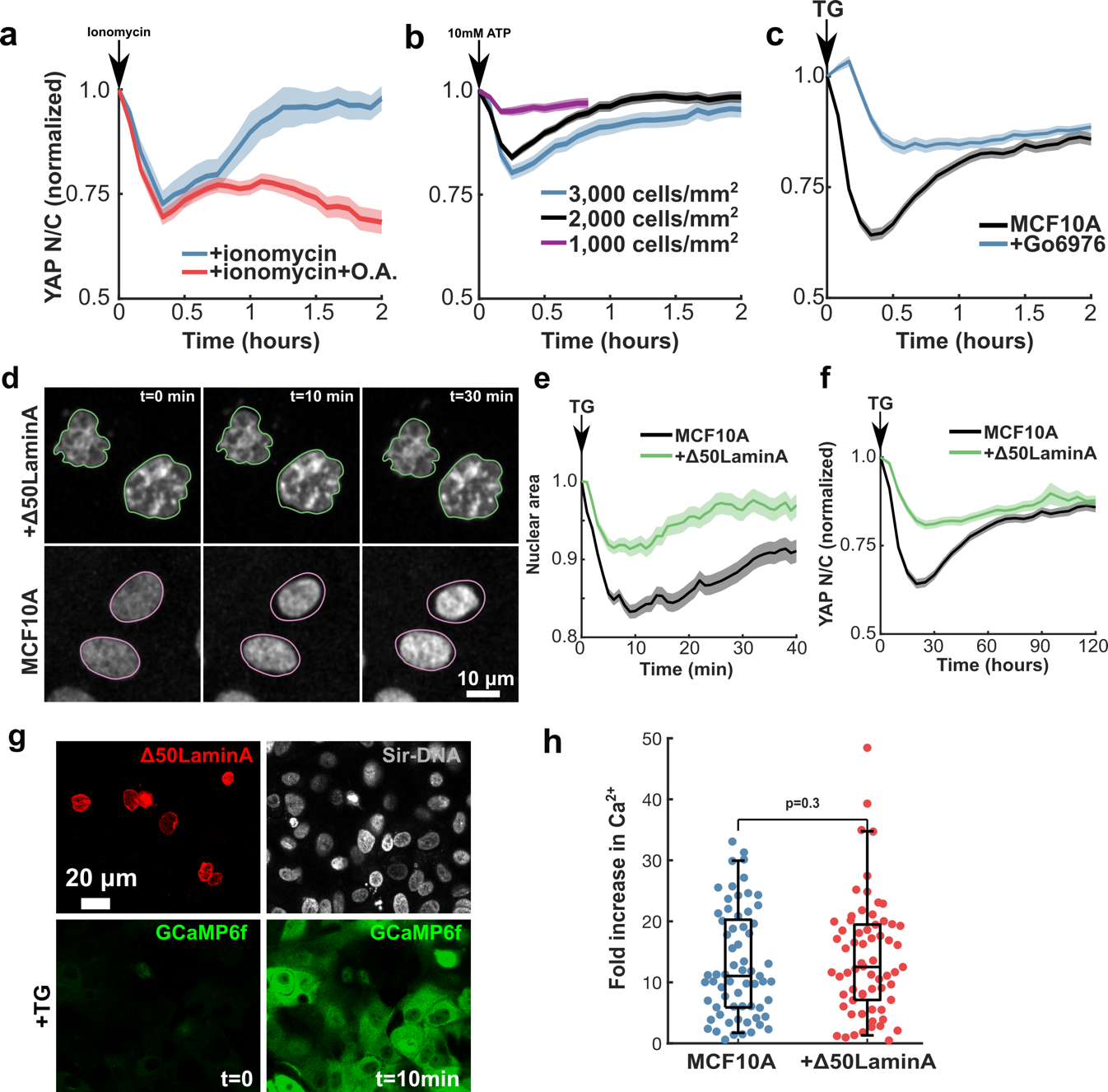

**Supplementary Fig. 3: Mechanism of YAP spatial reset.** (**a-c**) Mean YAP N/C for cells tracked over time under various conditions (All data are normalized to initial time point). (**a**) Treatment with ionomycin (N=47 tracks) or ionomycin and Okadaic acid (N=104 tracks). (**b**) Treatment with 10mM ATP at cell densities of 1,000 cells/mm^2^ (N=80 tracks), 2,000 cells/mm^2^ (N=162 tracks), and 3,000 cells/mm^2^ (N=186 tracks). (**c**) Go6976 pre-treatment + TG (N=104 tracks). Data in **a**-**c** are aggregated from two independent experiments for each condition. (**d**) Representative images of the nuclear compression following release of calcium with TG in MCF10A (left column) and MCF10A co-expressing ∆50LaminA (right column). (**e**) Normalized projected nuclear area for MCF10A cells (N=166 tracks), and MCF10A cells co-expressing ∆50LaminA (N=87 tracks) upon treatment with TG. Data in **b** and **c** are aggregated from two independent experiments. (**f**) Mean YAP N/C (normalized to initial time point) for MCF10A cells (N=137 tracks) and MCF10A cells co-expressing ∆50LaminA (N=106 tracks) upon treatment with TG. (**g**) Representative images of Thapsigargin-induced Ca^2+^ release. *Top:* Transient transfection of GCaMP6f cell line with ∆50LaminA-mCherry and cell nuclei stained with Sir-DNA. *Bottom*: GCaMP6f reporter with TG-treatment at t=0 and t=10 minutes. (**h**) Fold-increase in Ca^2+^ as measured by GCaMP6f signal at t=10 minutes normalized to mean signal at t=0. N_control_=65, N_Lamin_=65 cells. Box and whisker lines: black-center line is median value, box-bounds are inter-quartile range, and whiskers are 0.05 and 0.95 quantiles. P=0.3 from two-sided Mann-Whitney U-test (zval: -0.5262, ranksum: 4144). Repeated three times with similar results.

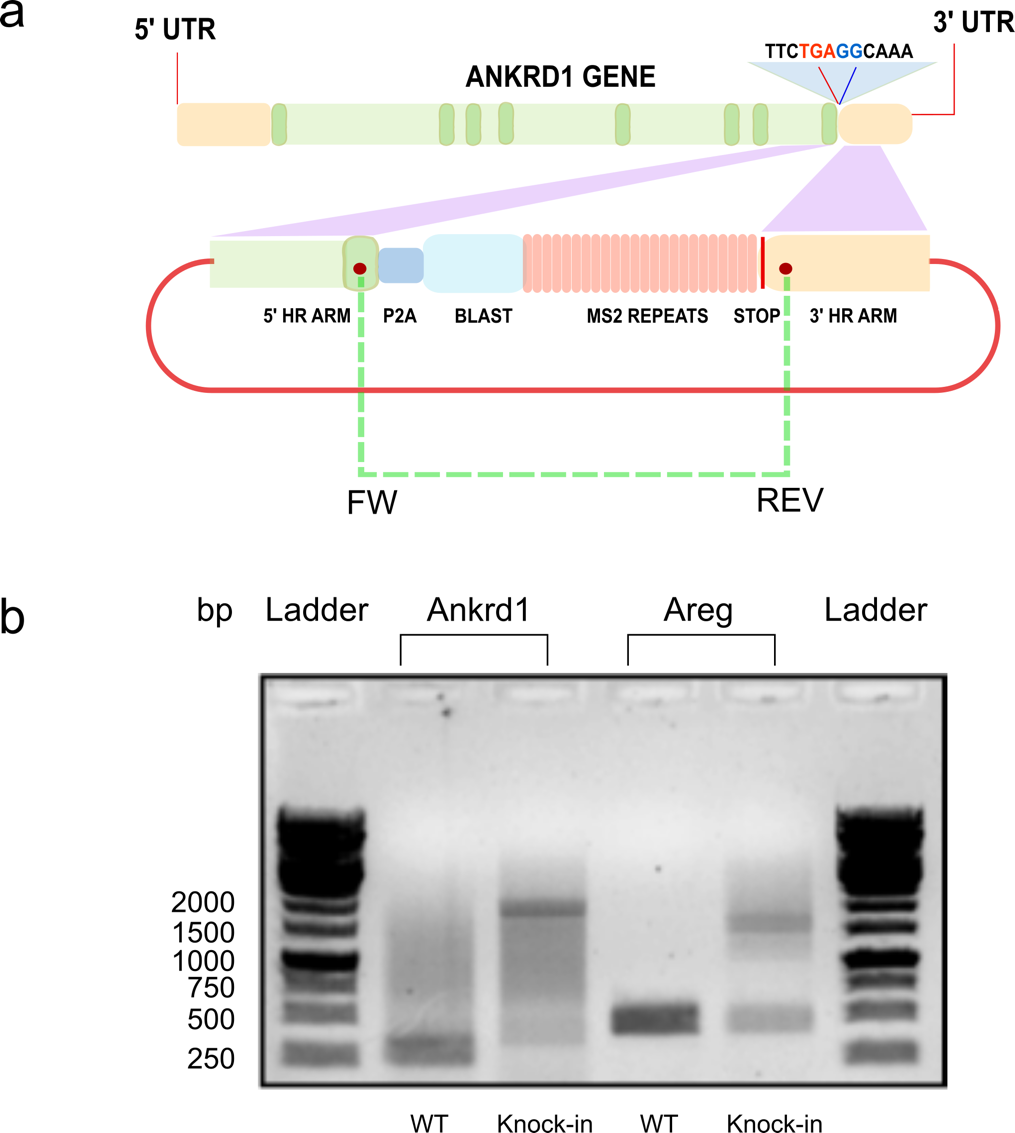

**­**

**Supplementary Fig 4: Validation of MS2 (transcriptional reporter cassette)-knockin cell lines. a.** Top: Cartoon of CRISPR-Cas9 based strategy for knocking in an MS2 transcription reporter cassette at the end of the coding sequence of ANKRD1 gene as shown in main text Figure 5a. Two sites in the 5’ and 3’ HR arms are marked to show locations of the FW and Rev primers used for PCR based validation of knock-in. Similar primer design was used to validate MS2 cassette knockin at the the end of the coding sequence of AREG gene.

**b.** Agarose gel electrophoresis of amplicons produced by PCR using Ankrd1 (lanes 2, 3) and AREG (lanes 4, 5) primers sets against total genomic DNA extracted from WT (wild type) and KI (knock-in) MCF10A cells.

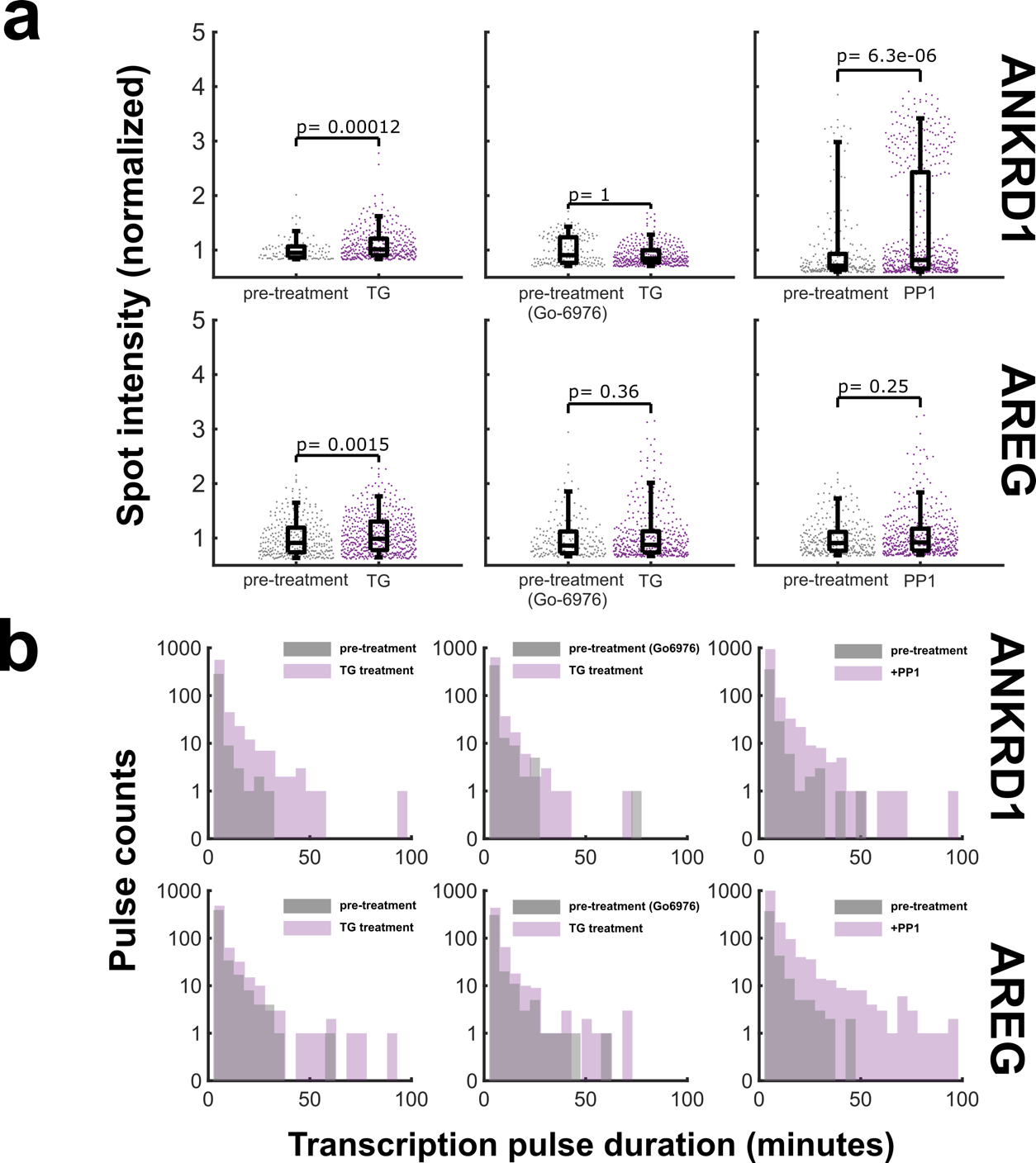

**Supplementary Fig. 5: Intensity and duration analysis of AREG and ANKRD1 gene transcription pulses under various conditions.** (**a**) Transcription spot intensity comparison between pre-treatment period (‘control’) and post-treatment period (“TG” or “PP1”). Spot intensity is the integrated fluorescence intensity pixel counts of each detected spot after subtracting local background signal. For each individual experiment, the transcription amplitude was normalized to the pre-treatment period such that the mean intensity is 1. The one-sided Mann-Whitney U-test P-value is reported for each pre-treatment: post-treatment comparison. *ANKRD1:* TG (P=1.2x10-4, Ranksum=20359, Z-val=-3.66, N_control_=105, N_TG_=368), +Go6976 (P=1.0, Ranksum=51121, Z-val=3.2, N_control_=168, N_TG_=375), +PP1 (P=6.3x10^-6^, Ranksum=74535, Z-val=-4.4, N_control_=229, N_PP1_=526). *AREG*: TG (P=0.0015, Ranksum=114063, Z-val=-2.95, N_control_=309, N_TG_=489), +Go6976 (P=0.36, Ranksum=56696, Z-val=-0.35, N_control_=191, N_TG_=409), +PP1 (P=0.25, Ranksum=114648, Z-val=-0.67, N_control_=297, N_PP1_=488). The distribution was generated by compiling data from all experiments from each condition. Box and whisker lines: black-center line is median value, box-bounds are inter-quartile range, and whiskers are 0.05 and 0.95 quantiles. (**b**) The distribution of pulse-durations are shown for each pre-treatment: post-treatment comparison, as in **a**. Pulse duration is defined as the time over which a nascent transcription spot was detectable and tracked.

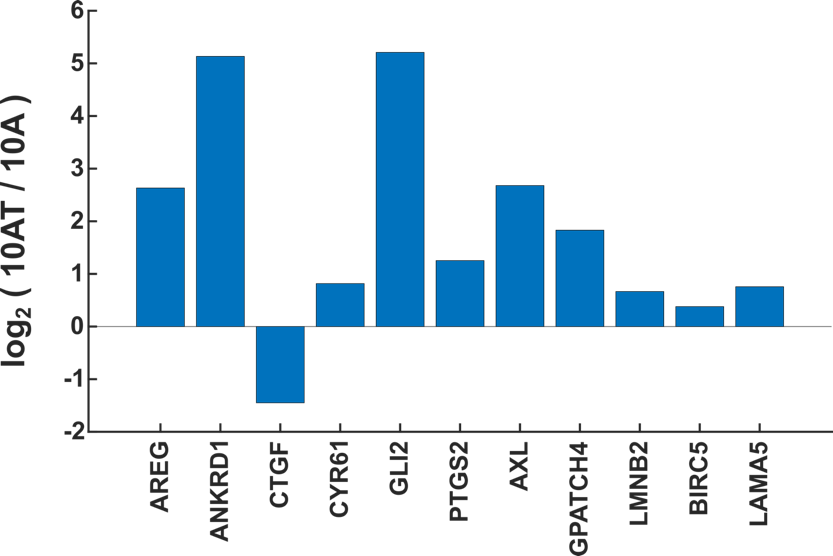

**Supplementary Fig. 6:** **Transformed mammary epithelial cells exhibit higher expression of YAP target gene.** Gene expression in Ras-transformed MCF10A (10AT) relative to MCF10A (10A) for genes known to be regulated by YAP activity based on reported RNA-seq measurements (See Methods)**.**

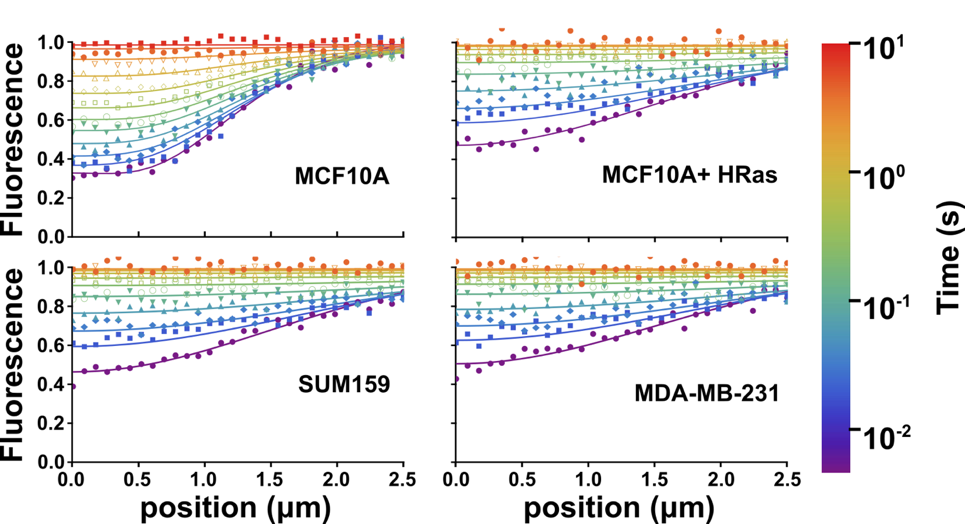

**Supplementary Fig. 7: Line FRAP reveals binding kinetics of YAP in normal and transformed breast epithelial cell lines.** A spatio-temporal FRAP model was used to fit high-resolution line FRAP experiments of YAP in MCF10A (N=30 cells), MCF10A+HRas (N=26 cells), SUM159 (N=30 cells), and MDA-MB-231 (N=30). Markers represent the measured mean fluorescence values as a function of time (color bar) and position from bleach center (x-axis). The model fit is shown in solid colored lines.

Supplementary Table 1: Line-FRAP model fitting. Fitted model parameter values for MCF10A, MCF10A + HRas, SUM159, MDA-MB-231 for either a pure-diffusion or reaction-diffusion model. Unrealistic values are highlighted in red. SSD is the sum of squared residuals between experiment data and model fit.

|  | Pure diffusion | | reaction-diffusion | | | | |
| --- | --- | --- | --- | --- | --- | --- | --- |
| cell line | D (μm^2^/s) | SSD | D_eff_ (μm^2^/s) | k_on_ (1/s) | k_off_ (1/s) | % Bound | SSD |
| MCF10A | 2.05±0.18 | 8.82±0.65 | 9.8±2.4 | 1.19±0.41 | 1.06±0.19 | 52±5 | 7.97±0.59 |
| MCF10A+HRas | 23.5±4.1 | 6.51±0.74 | 38.3±10.2 | 0.26±0.26 | 1.12±0.52 | 16±7 | 6.23±0.68 |
| SUM159 | 26.8±5.1 | 2.57±0.40 | 40.8±10.9 | 0.66±1.03 | 3.34±5.1 | 15±6 | 2.46±0.40 |
| MDA-MB-231 | 26.5±7.7 | 3.56±0.44 | 45.5±10.6 | 0.52±0.47 | 1.99±1.2 | 19±7 | 3.41±0.42 |

**Supplementary Movies List**

Supplementary Movie 1: 1.5 µM Latrunculin B treatment of MCF10A^YAP-GFP-KI^ cells.

Supplementary Movie 2: YAP localization vs. density in MCF10A^YAP-GFP-KI^ and HRas transformed MCF10A^YAP-GFP-KI^

Supplementary Movie 3: Highlighted MCF10A^YAP-GFP-KI^ cell with rapid changes of YAP localization.

Supplementary Movie 4: YAP N/C localization in MCF10A^YAP-GFP-KI^ cells, MCF10AT^YAP-GFP-KI^ cells and MCF10A^YAP-GFP-KI^ cells treated with the SRC inhibitor PP1.

Supplementary Movie 5: Serum-starved MCF10A^YAP-GFP-KI^ cells stimulated with serum-containing media.

Supplementary Movie 6: H1^YAP-GFP-KI^ hESC cells with a highlighted cell showing fluctuations in YAP localization.

Supplementary Movie 7: YAP localization dynamics at the wound edge.

Supplementary Movie 8: Thapsigargin-induced YAP localization in MCF10A control, Go6976 pre-treatment, and ∆50LaminA overexpression. Thapsigargin added before frame 1.

Supplementary Movie 9: Thapsigargin induced nuclear-deformation comparison between control and ∆50LaminA expressing cells.

Supplementary Movie 10: Tracking transcription kinetics of serum-starved MCF10A^AREG-MS2-KI^ cells before and after stimulation with serum-containing media.

Supplementary Movie 11: Accumulation of cytoplasmic ANKRD1 mRNAs in serum-starved MCF10A^Ankrd1-MS2-KI^ cells stimulated with serum-containing media.

Supplementary Movie 12: Tracking transcription kinetics before and after Thapsigargin treatment in MCF10A^ANKRD1-MS2-KI^ and MCF10A^AREG-MS2-KI^ cells.

Supplementary Movie 13: Transcription kinetics before and after mitosis in MCF10A^ANKRD1-MS2-KI^ and MCF10A^AREG-MS2-KI^ cells. The timestamp is with respect to cytokinesis.
